# Free fatty acids increase basal glucose uptake by adipocytes leveling it to insulin-stimulated uptake

**DOI:** 10.64898/2025.12.11.693656

**Authors:** Nikita V. Podkuychenko, Polina A. Bogdanova, Veronika S. Perelygina, Andrey N. Tikhonov, Gennady A. Badun, Maria V. Sudnitsyna, Alexander V. Vorotnikov

## Abstract

Increased free fatty acids (FFA) are considered a key factor in the development of insulin resistance in muscle and liver; however, their role in regulating glucose uptake in adipocytes remains controversial. Here, the effects of palmitate (PA) and oleate (OA) were studied in 3T3-L1 adipocytes with a focus on lipid accumulation and glucose uptake. We found that glucose rather than FFA availability promotes adipocyte maturation and fat accumulation associated with increased basal glucose uptake and altered expression of fatty acid oxidation markers (CPT-1 isoforms and UCP-1) toward lipid storage phenotype. Neither PA, nor OA altered insulin-stimulated glucose uptake by immature or mature adipocytes. Adipogenic differentiation of preadipocytes in the presence of rosiglitazone led to appearance of double effect of PA on glucose uptake by differentiated adipocytes, i.e. (1) PA dose-dependently increased basal glucose uptake, and (2) at high concentration PA suppressed insulin-stimulated glucose uptake. The effect of PA on basal glucose uptake was independent of mTORC1-mediated feedback in insulin signaling and persisted even when insulin signaling was inhibited by high-dose PA. Nonetheless, it was associated with GLUT4 exposure at the plasma membrane as reported by PA-induced, insulin-independent translocation of cMyc-GLUT4-mCherry chimera expressed in 3T3-L1 adipocytes. Thus, we conclude that only excessive FFA may context-dependently trigger classic insulin resistance in adipocytes, but otherwise FFA increase glucose uptake in adipocytes via GLUT4 mobilization.

## INTRODUCTION

Lipid metabolism disorders and insulin resistance are the hallmarks in pathophysiology of type 2 diabetes (T2D) (Accili et al., 2025, Reaven 1988). Insulin resistance is characterized by a reduced ability of insulin to stimulate glucose glucose clearance from the blood, leading to hyperglycemia and the manifestation of T2D (Roden, Shulman, 2019). Elevated levels of free fatty acids (FFA) in the blood plasma or inside cells critically contribute to the development of insulin resistance (Accili et al., 2025, Boden 2011, Shulman 2014). This phenomenon has been extensively studied in skeletal muscle and led to the concept of lipid-induced insulin resistance driven by increased FFA, which was later extended to hepatocytes (Petersen, Shulman, 2018, Samuel et al., 2010). However, adipocytes may not be susceptible to lipid-induced insulin resistance as these cells are adapted for storage and mobilization of large fat reserves. The mechanism of insulin resistance development in adipocytes is far from unraveled.

Palmitic (PA) and oleic acids (OA) are most abundant FFA in the body, however the saturated PA is more lipotoxic and detrimental with regard to insulin resistance. Thus, in cultured skeletal muscle cells (L6, C2C12) and rodent skeletal muscle fragments, PA inhibited insulin-stimulated glucose uptake (Alkhateeb et al., 2007, Den Hartogh et al., 2022, Dimopoulos et al., 2006, Gao et al., 2009, Hardy et al., 1991, Park et al., 2014), and Glut4 translocation (Hoehn et al., 2008), while it did not alter the basal glucose uptake. This effect of PA was reversed by unsaturated FFA (Gao et al., 2009, Park et al., 2014), which themselves increased glucose uptake (Fu et al., 2021, Park et al., 2014). Glucose uptake was reduced in muscle tissue fragments from animals following HFD (Hansen et al., 1998) or lipid infusion (Hoy et al., 2009), and also in humans after lipid infusion (Storgaard et al., 2004). It was also noted that AMPK activation may counteract the PA effects on skeletal muscle responses to insulin (Den Hartogh et al., 2022, Fu et al., 2021, Park et al., 2014).

Unlike skeletal muscle, the in vitro studies of FFA effects on glucose uptake by adipocytes yielded the contradictory results. In cultured model 3T3-L1 adipocytes (Fong et al., 1996, Grunfeld et al., 1981, Nugent et al., 2001, Tebbey et al., 1994, Usui et al., 1999) or in primary rat adipocytes (Haag et al., 2009, Hardy et al., 1991, Hebbachi, Saggerson, 2012, Hunnicutt et al., 1994, Joost, Steinfelder, 1985, Murer et al., 1992, Thode et al., 1989, Van Epps-Fung et al., 1997), different FFA variably increased basal glucose uptake (Den Hartogh et al., 2022, Fong et al., 1996, Hardy et al., 1991, Hebbachi, Saggerson, 2012, Hunnicutt et al., 1994, Joost, Steinfelder, 1985, Nugent et al., 2001, Tebbey et al., 1994, Thode et al., 1989, Van Epps-Fung et al., 1997, Usui et al., 1999). However, the FFA effects on insulin-stimulated glucose uptake were ambiguous. A number of studies reported that it is suppressed by FFA (Den Hartogh et al., 2022, Grunfeld et al., 1981, Hunnicutt et al., 1994, Tebbey et al., 1994, Van Epps-Fung et al., 1997), some studies found no effect (Fong et al., 1996, Joost, Steinfelder, 1985, Mukherjee et al. 1980, Murer et al., 1992), while other reported that FFA even enhance the uptake (Hardy et al., 1991, Hebbachi, Saggerson, 2012, Nugent et al., 2001, Thode et al., 1989, Usui et al., 1999). These inconsistencies could be related to methodological deviations in measuring the glucose uptake, including absence or different concentrations of glucose in the cell media, duration of cell exposure and different concentration of FFA, or use of different glucose tracers (2-DOG vs. ^14^C-U-glucose), etc. Thus, no definitive conclusion can be drawn on FFA effects in adipocytes, differences in saturated and unsaturated FFA action, their relation to glucose uptake and lipid-induced insulin resistance. An initial assumption that FFA may change the membrane fluidity (Grunfeld et al., 1981) was not confirmed (Nugent et al., 2001), but it was noted that FFA alter the lipid composition of the cell membrane, and lipoxygenase inhibitors counteract the effects of FFA on glucose transport in adipocytes (Nugent et al., 2001). The most detailed study suggested that FFA exert metabolic effects on glucose metabolism in adipocytes (Hebbachi, Saggerson, 2012) that are not consistent with Randle hypothesis of substrate competition for oxidation (Randle et al., 1963). Instead, the mechanism of FFA action in adipocytes was proposed to be related to increased requirement in glucose to generate 3-phosphoglycerate, which is needed for synthesis of glyceride glycerol and incorporation of external, re-esterified, or produced from glucose de novo FFA (Bally et al., 1960, Krycer et al., 2017, Krycer et al., 2020). It was also suggested that AMPK may also mediate glucose uptake in adipocytes, as it does in the muscle cells (Hebbachi, Saggerson, 2012), however, this has not been further investigated in depth. In adipose tissue biopsies from mice after HFD, basal glucose uptake was higher and not further increased by insulin (Talior et al., 2003), or hardly changed, and its stimulation by insulin decreased only after prolonged HFD (Hansson et al., 2018).

The action of insulin is mediated by insulin binding to its receptor and activation of an extensive insulin signaling network, including metabolic and mitogenic arms (Burchfield et al., 2025). Metabolic signaling is mediated by the insulin receptor substrate (IRS1) and subsequent activation of phosphatidylinositol-3-kinase (PI3K) and protein kinase B/Akt (Manning, Cantley, 2007, Petersen, Shulman, 2018). In turn, Akt phosphorylates TBC1D4 (*aka* AS160), which regulates the activity of Rab-GTPases responsible for translocation of GLUT4 from the cytosol to the outer cell membrane to enhance glucose uptake (Klip et al., 2019). Lipid infusion to healthy volunteers under hyperinsulinemic-euglycemic clamp conditions reduced glucose utilization from the bloodstream (Boden et al., 1994), suppressed insulin-stimulated skeletal muscle glucose uptake (Roden et al., 1996), and impaired PI3K activation (Dresner et al., 1999). This effects of FFA in muscle cells were attributed to intracellular accumulation of diacylglycerol, activation of nPKC, serine-directed phosphorylation and inhibition of IRS1 activity (Samuel et al., 2010). However, insulin signaling either was not impaired (Hoy et al., 2009, Storgaard et al., 2004) or was decreased in vitro only by high PA concentrations (Hoehn et al., 2008). It was suggested that reduced glucose utilization in skeletal muscle may not be due to impaired insulin signaling but rather due to metabolic alterations (Hoehn et al., 2008, Hoy et al., 2009, Storgaard et al., 2004). Strikingly, the effects of FFA of insulin signaling in adipocytes have not been ascertained.

Here, we investigated effects of excessive exogenous FFA on triglyceride accumulation in cultured 3T3-L1 adipocytes immediately and several weeks after their adipogenic differentiation, changes in the composition of carnitine palmitoyl transferase (CPT-1) isoforms and UCP-1 as the key markers of lipid metabolism, and glucose uptake by these adipocytes. We used rosiglitazone to accelerate maturation of adipocytes and traced the effect of palmitate on basal and insulin-dependent glucose transport in these cells. Having discovered that palmitate dose-dependently increases basal glucose uptake by adipocytes but suppresses insulin-dependent transport at high concentrations, we examined in detail the effect of palmitate on basal and insulin-dependent glucose transport into adipocytes. As a result, we found that palmitate stimulates basal glucose uptake by adipocytes independently of insulin cascade activation, and this effect of palmitate is mediated by an insulin-independent increase in the exposure of the glucose transporter GLUT4 at the adipocyte plasma membrane.

## MATERIALS AND METHODS

### Differentiation and culturing 3T3-L1 adipocytes

Culturing and differentiation of 3T3-L1 cells (ATCC, USA) were performed according to ATCC protocol or by essentially the same procedure, but with the presence of 2 µM rosiglitazone for 48 hours in differentiation medium (Zebisch et al., 2012). Briefly, the 3T3-L1 cells were grown to confluence in DMEM medium (PanEco, Russia) containing 4.5 g/l glucose, 1% L-glutamine (Thermo Fisher Scientific, USA), penicillin and streptomycin (P/S, 50 U/ml and 0.05 mg/ml; PanEco, Russia), and 10% newborn calf serum (NBCS; Intl Kang, China). Then cells were cultured for 48 hours in differentiation medium and then for 48 hours in post-differentiation medium. The differentiation medium was DMEM; 4.5 g/l glucose; 1% L-glutamine; P/S; isobutylmethylxanthine (IBMX, 0.5 mM; Sigma, USA); dexamethasone (Dex, 1 µM; Sigma, USA); insulin (Ins, 1 µg/ml; Sigma, USA); 10% fetal bovine serum (FBS, Cytiva, Germany). The post-differentiation medium was the same, but lacked IBMX and Dex. Successful differentiation was judged by formation of lipid droplets inside cells. Maintenance medium (DMEM, 4.5 g/l glucose; 1% L-glutamine; P/S; 10% FBS) was used for further culturing of adipocytes. The medium was exchanged every 48 hours. To obtain mature adipocytes, cells were cultured for 1.5 months post differentiation with the medium exchange every other day.

### Differentiation and culturing of ***С***2***С***12 myotubes

Mouse C2C12 myoblasts were cultured to a confluent monolayer in DMEM medium (PanEco, Russia) containing 4.5 g/L glucose, 1% L-glutamine (Thermo Fisher Scientific, USA), penicillin and streptomycin (P/S, 50 U/ml and 0.05 mg/ml; PanEco, Russia), and 10% fetal bovine serum (FBS, Cytiva, Germany). Cells were then cultured for 8–9 days in differentiation medium consisting of DMEM (4.5 g/L glucose, 1% L-glutamine, P/S) and 2% horse serum (HS). Successful differentiation was confirmed by the formation of large, elongated, multinucleated cells. The medium was exchanged every other day.

### FFA-albumin complexes and cell treatment

A 200 mM solution of sodium palmitate or sodium oleate (Sigma, USA) in 50% ethanol was pre-warmed to 60°C and added to a 20% BSA solution (BioSera, France) prepared in maintenance medium (DMEM, 4.5 g/l glucose) at a ratio of 1:25, followed by thorough mixing. This resulted in 8 mM stock solutions of BSA-PA or BSA-OA complexes and a final BSA:FFA ratio of 1:2.7 used in experiments. Cells were seeded in 12-well plates. Cells were incubated with 1 mM palmitate or 1 mM OA complexed with BSA for 48 hours. As a first control, cells incubated for 48 hours in the presence of 2.5% (0.38 mM) BSA solution (BioSera, France) were used. As a second control, cells were incubated for 48 hours in the maintenance medium only.

### Measurement of triglyceride accumulation in adipocytes

The formation of lipid droplets and the increase in triglyceride content in 3T3-L1 adipocytes before and after culturing in the presence of palmitic or oleic acid complexed with BSA were assessed using fluorescence microscopy. Live cells were mixed with fluorescent dyes BodiPy-FL (triglyceride staining, absorption/emission maxima 502/511 nm; dilution 1:1000; Thermo Fisher Scientific, USA) and Hoechst 33342 (nuclear DNA staining, absorption/emission maxima 346/460 nm; dilution 1:1000). Then cells were incubated for 10-15 min at 37°C. Confocal imaging was performed using Leica Stellaris 5 microscope. The obtained images were processed using ImageJ software (NIH, USA). The number of cells was counted by stained nuclei, and the average fluorescence area occupied by lipid droplets was calculated per cell to estimate lipid droplet size as a proxy of triglyceride accumulation. Microsoft Excel was used for statistical analysis to obtain mean ± S.D. values.

### Stable 3T3-L1 cell line expressing Myc-Glut4-mCherry

Stable 3T3-L1 cell line expressing Myc-Glut4-mCherry fusion protein (Lim et al., 2015) was established by lentiviral transduction followed by fluorescence-activated cell sorting (FACS). Lentiviral particles were produced in HEK293T cells via calcium phosphate transfection using a three-plasmid system: the transfer plasmid pLenti-Myc-Glut4-mCherry (Addgene #64049), the packaging plasmid pCMV-dR8.2 (Addgene #8455), and the envelope plasmid pMD2.G encoding VSV-G (Addgene #12259). Conditioned medium was collected 48 hours post-transfection, clarified by centrifugation at 1000 g for 10 minutes, and used to transduce undifferentiated 3T3-L1 fibroblasts (ATCC, USA) for 24 hours in the medium supplemented with Polybrene (5 µg/ml, Sigma, USA) and the viral supernatant diluted 1:2 with fresh growth medium. The transfection and transduction efficiencies were assessed by mCherry fluorescence.

Transduced cells were selected by FACS on a BD FACSAria™ III flow cytometer (BD, USA). Cells were cultured to 70% confluence and prepared as a single-cell suspension at concentration of 10 cells/ml. Cell sorting was performed in the Purity mode using an 85 µm nozzle. The mCherry signal was detected using a 488 nm excitation laser with an emission filter set to 550–650 nm. Non-transduced 3T3-L1 fibroblasts served as a negative control for gating. The sorted population exhibiting medium mCherry fluorescence was collected in culture medium for further expansion and experiments.

### GLUT4 transporter localization by confocal microscopy

3T3-L1 cells stably expressing the Myc-Glut4-mCherry fusion protein were differentiated into mature adipocytes using standard protocol recommended by ATCC. The differentiated cells were cultured in DMEM-HG medium with 10% FBA for 1 week and then incubated for 48 hours in the fresh medium with addition of either BSA (2.5%) or 1 mM PA complexed with BSA. Where appropriate, cells were stimulated with 100 nM insulin for 15 minutes prior to confocal imaging.

For immunocytochemical staining, cells were washed twice with PBS and fixed with 4% formaldehyde for 10 minutes. After fixation, the cells were stained for nuclei with Hoechst 33342 (Invitrogen, H3570, USA), washed three times with PBS, and stored at +4°C in PBS containing 0.09% sodium azide (NaN). To visualize the chimera exposed on plasma membrane, c-Myc antibody (Invitrogen, #PA5-85185, USA) and secondary antibody conjugated with Alexa Fluor 488 (Invitrogen, #A-11034, USA) were used. Leica Stellaris 5 confocal laser scanning microscope (Leica Microsystems, Germany) was used to acquire fluorescence. The images were acquired using a 20x long-distance plan-apochromatic objective. Alexa Fluor 488, mCherry, and DAPI fluorescence were excited by an optical (fiber) IR laser at wavelengths of 488 nm, 543 nm, and 360 nm, respectively, with emission detected in the ranges of 500–550 nm, 570–620 nm, and 450–470 nm. Series of optical sections (Z-stacks) were obtained with a 0.5 µm step. Image processing and analysis (deconvolution, projection construction) were performed using LAS X software (Leica Microsystems) and ImageJ (NIH, USA).

### [^3^H]-2-deoxyglucose uptake

Cells pre-incubated with PA or OA were washed and placed in glucose-free DMEM medium for 10 min. Then, some cells were stimulated with 100 nM insulin for 15 minutes. Subsequently, 100 µM 2-deoxyglucose (2-DOG) and 0.1 Ci/mol [3H]-2-deoxyglucose (USA, ARC) as a tracer were added to the cells for 10 minutes. The reaction was stopped by adding cold Hanks’ solution (PanEco, Russia) containing 4.5 g/l glucose. Immediately after this, cells were frozen. Cells were detached from the plates by adding RIPA lysis buffer. Samples were homogenized by passing through an insulin syringe. Samples were centrifuged at 13,000g for 10 minutes, the supernatant was transferred to scintillation vials with Ultima Gold scintillation fluid (PerkinElmer, USA), mixed, and the β-radiation of 2-DOG was measured using a RackBeta counter (USA).

### Western blotting

After the experiment, cells were lysed in RIPA buffer with phosphatase and protease inhibitors (Thermo scientific, USA). The resulting cell lysates were passed several times through an insulin syringe to disrupt large cell fragments and DNA. Cell lysates were then centrifuged for 10 minutes at 10,000 g and 4°C. The supernatant was carefully collected, avoiding the pellet and floating lipids. The volume was then equalized in all samples. After adding 4x Laemmli sample buffer, samples were incubated at 90°C for 20 minutes. Prepared samples were stored at -20°C. Electrophoretic separation of proteins was performed by the Laemmli method (Laemmli, 1970) in the presence of sodium dodecyl sulfate (SDS). Polyacrylamide gels (4% stacking and 10% separating) were used, as well as commercial 4-12% gradient gels (BioRad, USA) and a set of prestained protein markers (BioRad, USA) with known molecular weights. A Criterion Cell system (BioRad, USA) was used for electrophoresis. Protein separation was performed at 15 mA and 30 mA per gel plate for stacking and separating gels, respectively. After protein separation, electrotransfer was performed. Completeness of transfer was monitored by staining the final gels with Coomassie R-250. A Criterion Cell system (BioRad, USA) was used for protein electrotransfer. A wet transfer protocol from BioRad was used to transfer proteins onto PVDF membranes (Merck, Ireland). Electrotransfer was carried out for 3 hours at 300 mA. The membrane was blocked with a 5% solution of skim milk (AppliChem, Spain) in TBST for one hour. Membranes were then washed three times with TBST for 10 minutes each and incubated with primary antibodies in a solution of 0.1% BSA and 0.1% NaN3 in TBS at the dilution recommended by the manufacturer overnight at 4°C. The next day, the membrane was washed 3 times for 10 minutes with TBST and incubated with horseradish peroxidase-conjugated secondary antibodies in a 1% skim milk solution at the recommended dilution at room temperature for one hour. Proteins were visualized using enhanced chemiluminescence (ECL) with Clarity™ Western ECL Substrate (1705061, BioRad, USA). Chemiluminescence was detected using a gel documentation system Infinity 3000 WL/LC (Vilber Lourmat, France). The obtained results were processed in ImageJ software. Sample loading was normalized to vinculin staining. The following primary antibodies were used for western blotting: anti-CPT1A (#15184, Proteintech, USA), anti-CPT1B (#22170, Proteintech, USA), anti-SCD1 (#2438, CellSignaling, USA), anti-UCP1 (#14670, CellSignaling, USA), and anti-vinculin (#9131, Sigma, USA). As secondary antibodies, goat anti-rabbit IgG (#ab6721, Abcam, USA) or goat anti-mouse IgG (#ab97023, Abcam, USA) conjugated with horseradish peroxidase were used.

### Statistical Analysis

The data are presented as means ± SD of at least two independent experiments. The *p* values were calculated using Student’s t test or one-way ANOVA and specified in the figure legends. Differences were considered statistically significant at *p* ≤ 0.05.

## RESULTS

Typically, glucose uptake by adipocytes is studied within 1-4 days after complete differentiation, which requires 10 days for 3T3-L1 cells (Rubin et al., 1978, Zebisch et al., 2012). However, these adipocytes are usually multilocular, and their metabolism and its regulation may further change as adipocytes mature and acquire a unilocular phenotype (Berger, Géloe n, 2022, Kim et al., 2019, Salans et al., 1968, Ye et al., 2022, Yu, Li, 2017). It has been even suggested that adipocyte size rather than obesity itself predicts T2D independent of insulin resistance (Weyer et al., 2000). Therefore, we explored whether the model 3T3-L1 adipocytes accumulate fat in the long-term cultures, alter lipid droplet morphology and glucose uptake. Also guided by common appreciation that excess FFA may contribute to insulin resistance, at least in muscle cells, we extended these studies to explore how excessive PA and OA, the two major FFA in organism, affect lipid droplet morphology and glucose uptake in 3T3-L1 adipocytes.

### Exogenous palmitate or oleate do not alter lipid droplet morphology and glucose uptake in 3T3-L1 adipocytes

Fig. 1A shows that lipid droplet morphology is dramatically different in 3T3-L1 adipocytes immediately after the standard differentiation or after their subsequent culturing for 45 days in serum and high-glucose containing medium. Whereas “young” adipocytes have small lipid droplets of multilocular appearance immediately after differentiation, the adipocytes grown for up to 1.5 months exhibit “mature” phenotype with mostly unilocular lipid droplets. Consistently, the area occupied by lipid droplets in immature cells (Fig. 1B) is on average about 10-times larger than that in mature adipocytes (Fig. 1C). This demonstrates that over time adipocytes actively accumulate fat and mature by changing morphology. They use external glucose as a source of triglycerides by converting it to both glyceride glycerol, and to fatty acids as long as the availability of glucose is more than 1 mM (Bally et al., 1960, Krycer et al., 2017, Krycer et al., 2020, Saggerson, 1972).

**Fig. 1.**
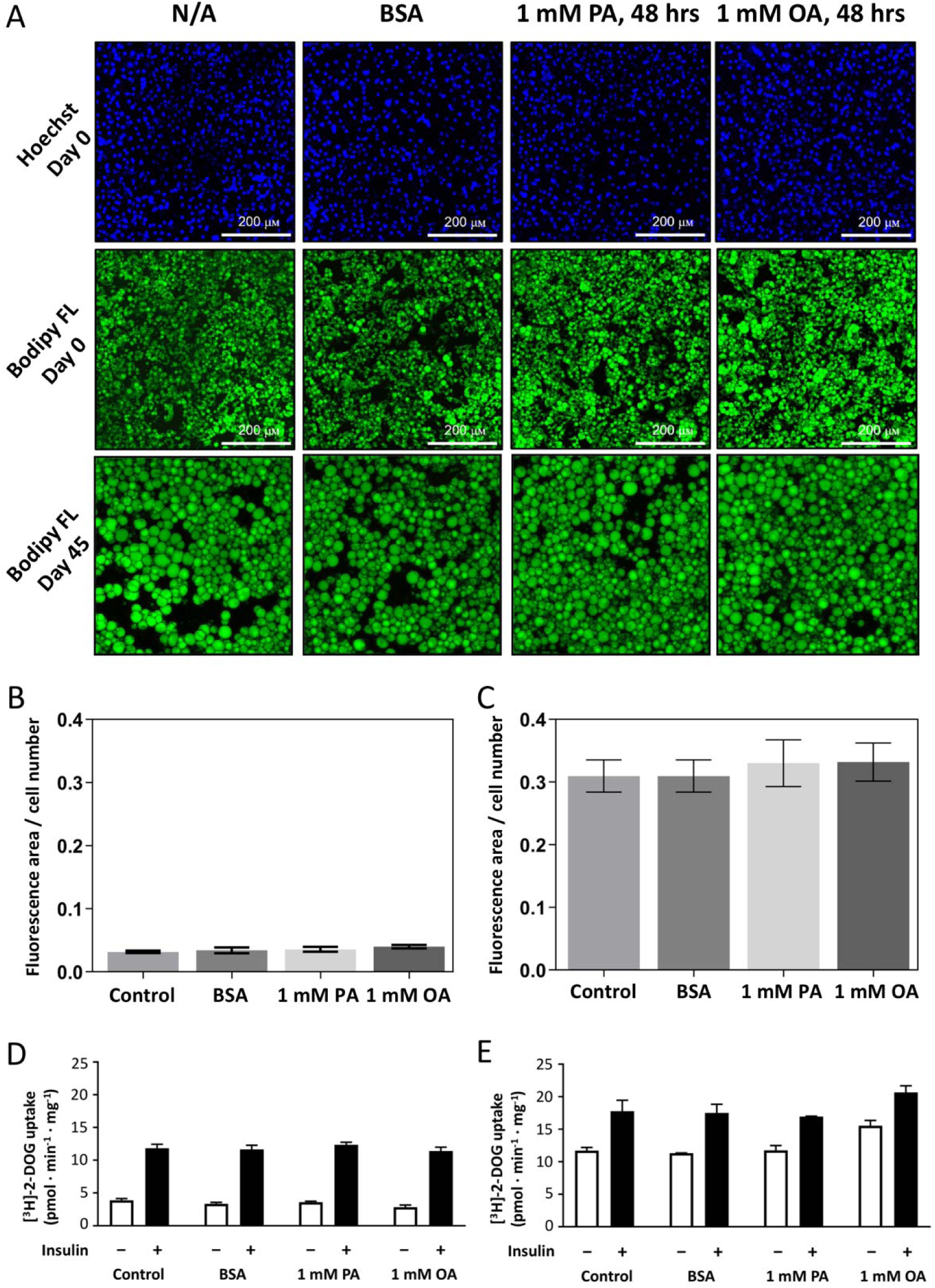
FFA treatment does not affect lipid droplet area or glucose uptake in 3T3-L1 adipocytes. **(A)** Confocal images of adipocytes stained with Bodipy FL immediately after completion of standard differentiation for 10 days (Day 0), or after 45 days in culture with DMEM-HG and 10% FBS (Day 45). Two days before the analysis, the cell medium was exchanged for that without (N/A), or with 2.5% defatted BSA (BSA), 2.5% BSA complexed with palmitate (1 mM PA), or oleate (1 mM OA) as indicated. After 48 hours, cells were washed and stained for lipid droplets (Bodipy FL) or nuclei (Hoechst) as indicated. **(B)** and **(C)** Bodipy FL fluorescence area in lipid droplets immediately after adipocyte differentiation (B, day 0) or 45 days after differentiation C, day 45), normalized to cell number. Data are presented as mean ± SD from 4-5 fields of view (n=3). **(D)** and **(E)** Basal (white columns) and insulin-stimulated (black columns) [^3^H]-2-DOG uptake by adipocytes immediately after (D) or 45 days (E) after differentiation. Data are presented as mean ± SD from 2 experiments performed in triplicates.

Aided by insulin, the external FFA delivered by albumin also easily incorporate into triglyceride as evidenced by their postprandial uptake. However, whether adipocytes can easily accumulate the external FFA in the absence of insulin and at relatively low ambient level of FFA is not obvious. To address this question, we additionally treated the immature and mature adipocytes by complexes of PA or OA with albumin. We treated adipocytes for 2 days with 1 mM of these FFA in their ratio to albumin of 2.7, assuming that their normal systemic levels are in the range of 0.5-0.6 mM (Arner, Ryden, 2015, Karpe et al., 2011) and there are 3 high affinity FFA binding sites in albumin (Curry 2003). We reasoned that adipocytes would experience an increased exposure to external FFA and enlarge their lipid droplets by converting them to triglycerides. However, this apparently was not the case, as evidenced by the lack of increase in the area occupied by lipid droplets both in the “young”, and mature adipocytes (Fig. 1A-C). We therefore conclude that both immature, and mature adipocytes effectively exchange FFA with the outer space via constant lipolysis and esterification, the essential feature of the fat cells.

To further explore if the excessive external FFA may induce insulin resistance in adipocytes, we assessed glucose uptake using [^3^H]-2-deoxyglucose ([^3^H]-2-DOG) as a tracer. We did observe any FFA-induced reduction in glucose uptake by both types of the adipocytes (Fig. 1D,E). Moreover, mature adipocytes revealed even greater 2-DOG uptake in the presence of insulin. However, most surprisingly, the mature adipocytes had a more that 3-fold higher basal glucose uptake, which accounted for its less fold activation by insulin (on average, 1.5-fold vs. 3.5-fold in immature “young” adipocytes). Altogether, these results argue that mature unilocular adipocytes have increased glucose uptake unaltered by excessive exogenous FFA present throughout 2 days in the cell medium. This is consistent with notion that adipocytes constantly turnover their triglycerides, which would require more glucose supply to maintain the activity along fat accumulation (Morigny et al., 2021).

### Adipocytes alter expression of FFA oxidation markers upon maturation

Next, we asked if the increased triglyceride content in mature adipocytes is associated with altered expression of the enzymes that govern FFA oxidation. CPT-1 is a bottleneck for transfer of fatty acids to mitochondria for oxidation (McGarry et al., 1977, Newgard 2018); it has been implicated in intracellular routing of fatty acids and dysregulated metabolism leading to T2D (McGarry 2002). The low expression of CPT-1 in white adipose tissue, as compared to brown fat and other tissues (Warfel et al., 2017), suggests that adipocyte maturation is associated with re-routing of fatty acids from mitochondria to storage compartment. Similarly, UCP-1 content is a marker of mitochondrial oxidation activity (Rosen, Spiegelman, 2014) and may indirectly reflect intensity of fatty acid metabolism.

We used western blotting to assess the content of CPT-1 isoforms and UCP-1 in immature and mature 3T3-L1 adipocytes (Fig. 2A,B). The content of CPT-1A, a ubiquitous CPT-1 isoform across tissues, was higher in preadipocytes, reduced in post-differentiated adipocytes, and further decreased in mature adipocytes after 45 days in cell culture. Strikingly, immature adipocytes express large amounts of CPT-1B isoform, which is abundant in muscle and brown fat cells (Fig. 2A,B). They also express UCP-1, indicative of their high fatty acid oxidation capacity. In contrast, mature adipocyte lacked both CPT-1B, and UCP-1, which is consistent with their little fatty acid oxidation and increased fat storage in large lipid droplets, which accumulate over time (cf. Fig. 1A,C). To extend these findings, we compared the CPT-1 and UCP-1 content in skeletal muscle C2C12 myotubes. Thus, we differentiated C2C12 myoblasts and found that many, although not all, cells fused into myotubes as evident by their multi-nuclei fiber-like morphology and skeletal troponin I staining (Fig. 2C). This cells contained both CPT-1A and CPT-1B in amounts comparable to those in preadipocytes and immature adipocytes, respectively (Fig. 2A,B). They also contained UCP-1, though in less and variable amounts than in immature adipocytes, which is probably due to irregular and incomplete differentiation of these cells as noted above. Altogether, these results suggest that adipocytes dramatically alter their lipid metabolism during maturation and acquire pronounced lipid storage phenotype and increased glucose utilization.

**Fig. 2.**
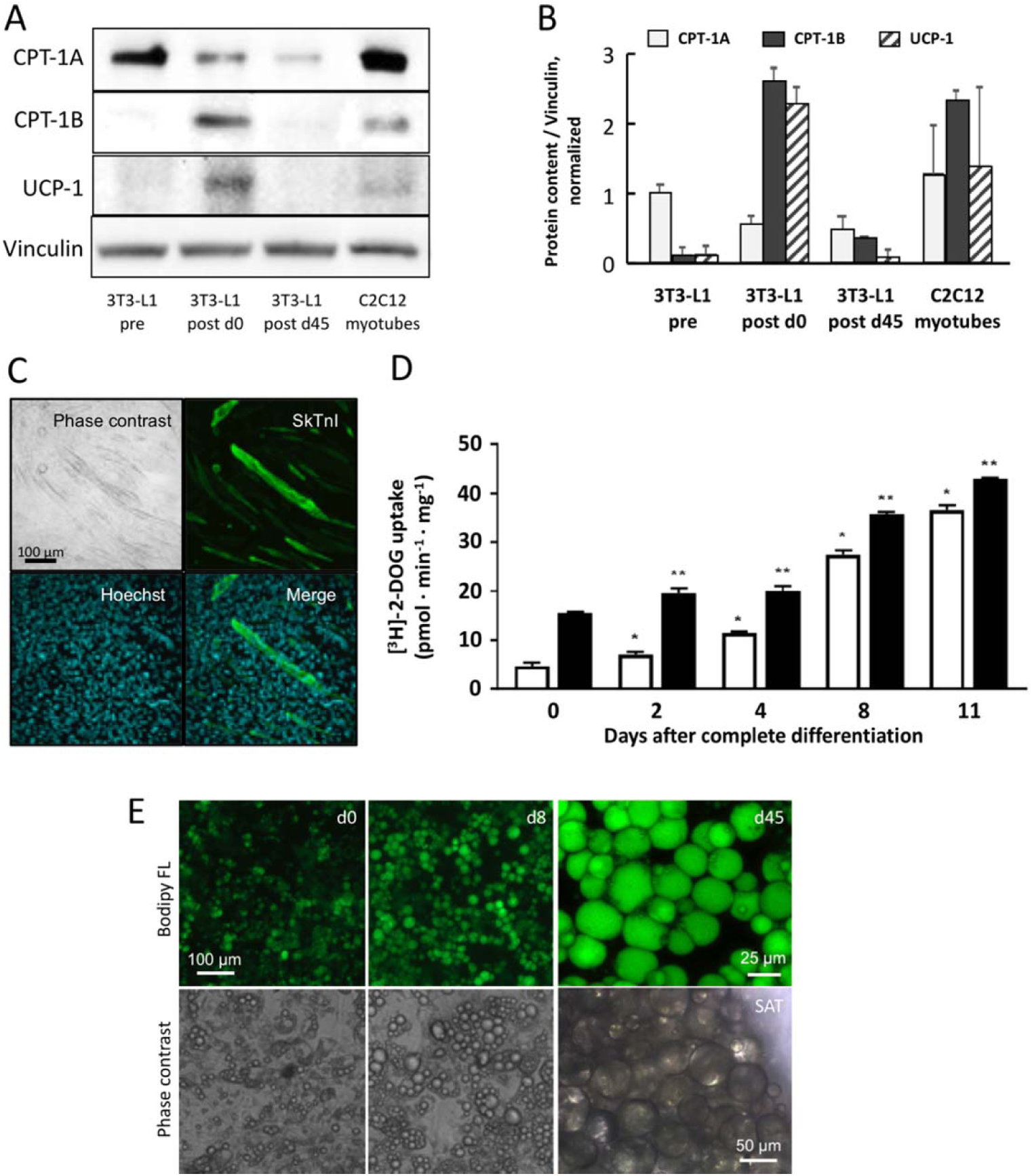
3T3-L1 adipocytes alter expression of CPT-1 and UCP-1 driven by increased glucose availability during maturation. Representative western blots **(A)** and their quantitation **(B)** show differences in the content of CPT-1A, CPT-1B, and UCP-1 in preadipocytes (3T3-L1 pre), newly differentiated adipocytes (3T3-L1 post d0), mature adipocytes (3T3-L1 post d45), and C2C12 myotubes shown for reference. **(C)** The C2C12 myotubes were visualized by phase contrast, stained for skeletal troponin I (skTnI) and for nuclei (Hoechst) to assess effectiveness of myogenic differentiation. **(D)** Confocal images of adipocytes stained with Bodipy FL for lipid droplets (upper row) or visualized by phase contrast (lower row) just after differentiation (d0), and subsequent culturing in high glucose for 8 days (d8) and 45 days (d45). The lower right image shows human adipocytes at the edge of subcutaneous adipose tissue biopsy (SAT) taken from an obese individual. Scale bars are given for comparison of cell sizes. (E) Temporal dynamics of basal and insulin-stimulated glucose uptake during the early adipocyte maturation measured with [^3^H]-2-DOG as a tracer at days indicated on the abscissa. Data are shown as means ± SD from 2 experiments in triplicates; *, p <0.05; **, p <0.01.

### Adipocyte maturation is associated with increased glucose demand

The observed differences between immature and mature adipocytes suggest that newly differentiated 3T3-L1 adipocytes are more beige-like and gradually transform into white adipocytes due to increased glucose availability in cell culture medium. To explore this hypothesis, we closely traced changes in glucose uptake over first several days post differentiation. Adipocytes successively increased both basal and insulin-stimulated glucose uptake, yet the former was quantitively more pronounced resulting in diminution on fold stimulation by insulin (Fig. 2D). Basal glucose uptake increased up to 7-fold, but the insulin-stimulated uptake increased hardly more than twice by day 8 after the end of differentiation, thus resulting in about 3-fold decrease in fold stimulation by insulin. Thereafter glucose uptake tended to plateau, which suggests that cells might have reached maximum response and exhausted their glucose transport machinery. Notably, while the decrease in fold stimulation by insulin formally depicts loss of insulin action, it is largely due to increase in the basal uptake rather than loss of the insulin-stimulated response, as would be expected from classic insulin resistance in skeletal muscle.

The lipid droplet morphology also markedly changed by day 8 post differentiation upon cell culturing in high glucose medium (Fig. 2E, d8 vs. d0). While initially multilocular, adipocytes adopted the unilocular phenotype, apparently increased in size and lipid content. When the medium was refilled every other day until day 45, the cells attained clearly unilocular phenotype with one large lipid droplet virtually in most cell (Fig. 2E, d45). Notably, such an appearance is typical of human adipocytes as evidenced by the light microscopy imaging of subcutaneous fat biopsy from an obese individual (Fig. 2E, the lower image on the right). Yet, human cells appear twice as large as mature rodent 3T3-L1 adipocytes, which may reflect specie-specific differences, insufficient 3T3-L1 adipocyte maturation, or both.

### Rosiglitazone during adipogenic differentiation confers PA-sensitivity to glucose uptake

As the early changes in glucose uptake were closely associated with changes in lipid droplet morphology, we hypothesized that forced stimulation of lipid anabolic program may accelerate the development of increased glucose uptake. Thus, we introduced rosiglitazone for two critical days of adipogenic differentiation of 3T3-L1 preadipocytes as described (Zebisch et al., 2012). The cells differentiated for 10 days by this procedure had slightly increased basal glucose uptake than adipocytes differentiated in the absence of rosiglitazone, and their glucose uptake was similarly stimulated by insulin 4-5-fold (Fig. 3A). In addition, both cells expressed similar levels of CPT-1 isoforms and UCP-1 as assessed on day 2 post differentiation (Fig. 3A).

**Fig. 3.**
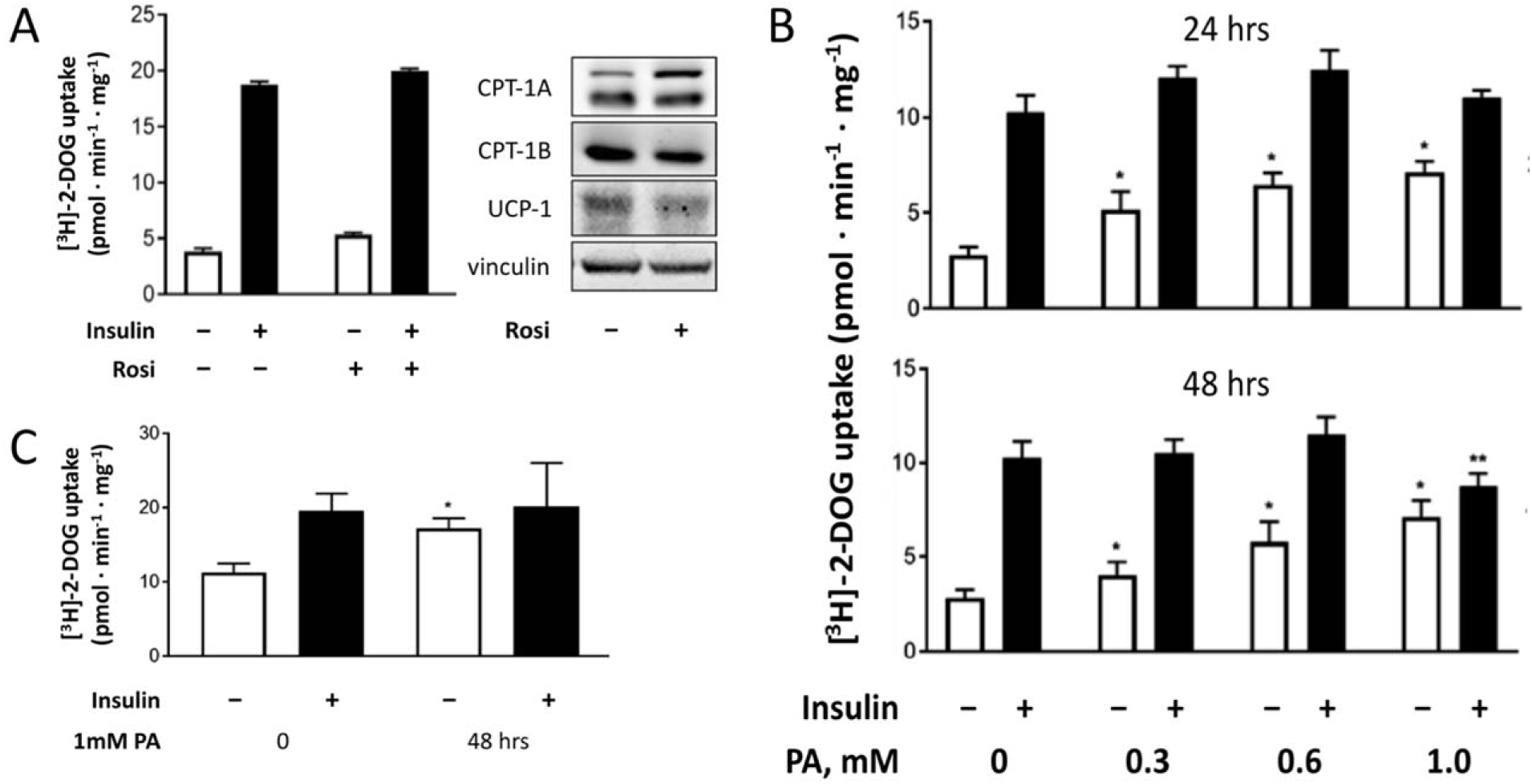
Rosiglitazone-assisted differentiation of 3T3-L1 adipocytes confers PA sensitivity to glucose uptake. **(A)** Comparison of glucose uptake (*left*) and the content of CPT-1 isoforms and UCP-1 (*right*) in adipocytes 2 days after complete adipogenic differentiation conducted in the absence or presence of rosiglitazone (Rosi) for 2 days in differentiation cocktail as indicated. **(B)** Dose-dependence of palmitate (PA) effect on glucose uptake by adipocytes differentiated in the presence of rosiglitazone and treated either with BSA alone (2.5%) or with PA-BSA complexes (2.7:1 ratio) for 24 hours (*upper panel*) or 48 hours (*lower panel*). **(C)** Glucose uptake of human adipocytes differentiated from ADSC in the presence of rosiglitazone treated with BSA (2.5%, 0 mM PA) or 1 mM PA in complex with BSA (2.7:1 ratio) for 48 hours. Data are shown as means ± SD from n≥ experiments performed in triplicates; *, p <0.05 compared to uptake in the absence of PA and insulin; **, p <0.01 compared to that in the absence of PA.

The rosiglitazone-treated cells displayed different sensitivity of glucose uptake to PA treatment than those differentiated in the absence of rosiglitazone. While in the latter adipocytes neither PA, nor OA altered basal or insulin-stimulated glucose uptake (Fig. 1D), PA dramatically and dose-dependently increased basal glucose uptake in rosiglitazone-treated 3T3-L1 adipocytes as early as at day 0 post differentiation (Fig. 3B). This increase in basal glucose uptake was comparable in cells treated by PA for 24 or 48 hours, whereas it was only marginally increased or not affected by insulin in adipocytes treated by PA for 24 and 48 hours, respectively. However, at high concentration (1 mM) PA significantly decreased insulin-stimulated glucose uptake as compared to PA untreated cells, or those treated with lower concentrations of PA (Fig. 3B). Altogether, these results suggest that rosiglitazone presence during adipogenic differentiation produces minimal early changes in fatty acid utilization pathways and glucose uptake, but confers sensitivity of glucose uptake to external PA.

To further substantiate these results, we isolated human ADSC from subcutaneous fat of obese subjects (Stafeev et al., 2019). As these cells may display lower proliferative and adipogenic potential, they were differentiated for 10 days by the same protocol with rosiglitazone (Zebisch et al., 2012).

Strikingly, when treated by 1 mM PA for 48 hours, these adipocytes displayed ∼1.6-fold higher basal glucose uptake while their response to insulin was not altered (Fig. 3C). This further confirms that excessive exogenous PA primarily increases basal glucose uptake by adipocytes.

### Increased basal glucose uptake is spared from mTORC1-mediated feedback in insulin signaling

By far, our observations indicated that PA increases basal glucose uptake by adipocytes, while in high doses and/or with prolonged action it may also blunt insulin-stimulated glucose uptake, the hallmark of insulin resistance (Fig. 3B,C). The mTORC1-mediated feedback to IRS-1 has been implicated in both inhibition (Burchfield et al., 2025, Samuel et al., 2010), and activation (Brannmark et al., 2013) of insulin signaling to glucose uptake. To address the possible involvement of the feedback to PA effects on glucose uptake we used rapamycin to block the mTORC1-mediated feedback in 3T3-L1 adipocytes differentiated in the presence of rosiglitazone. To distinguish between two possibly different actions of PA on basal and insulin-stimulated uptake, we used different PA concentrations and duration of treatment of freshly differentiated 3T3-L1 adipocytes.

We first confirmed that mTORC1-mediated feedback is operative in our cells and can be abolished by rapamycin. Rapamycin dose-dependently inhibited insulin-induced phosphorylation of S6K1, a reporter substrate of mTORC1, and this was paralleled by increased phosphorylation of Akt and AS160, the key mediators of insulin signaling branch to glucose uptake (Fig. 4A). The S6K1 response to insulin was completely ceased by 100 nM rapamycin. Treatment of adipocytes by 0.5 mM PA for 24 hours resulted in the comparably increase in both basal, and insulin-stimulated phosphorylation of S6K1 (Fig. 4B), suggesting that mTORC1is activated following the cell treatment by PA. Howe er, PA also upregulated phosphorylation of Akt and AS160, and 100 nM rapamycin failed to decrease this PA-induced response of Akt and AS160 (Fig. 4C). Taken together, these results indicate that upregulation of insulin signaling to glucose uptake by PA is independent of PA signaling to mTORC1 or mTORC1-mediated feedback in 3T3-L1 adipocytes.

**Fig. 4.**
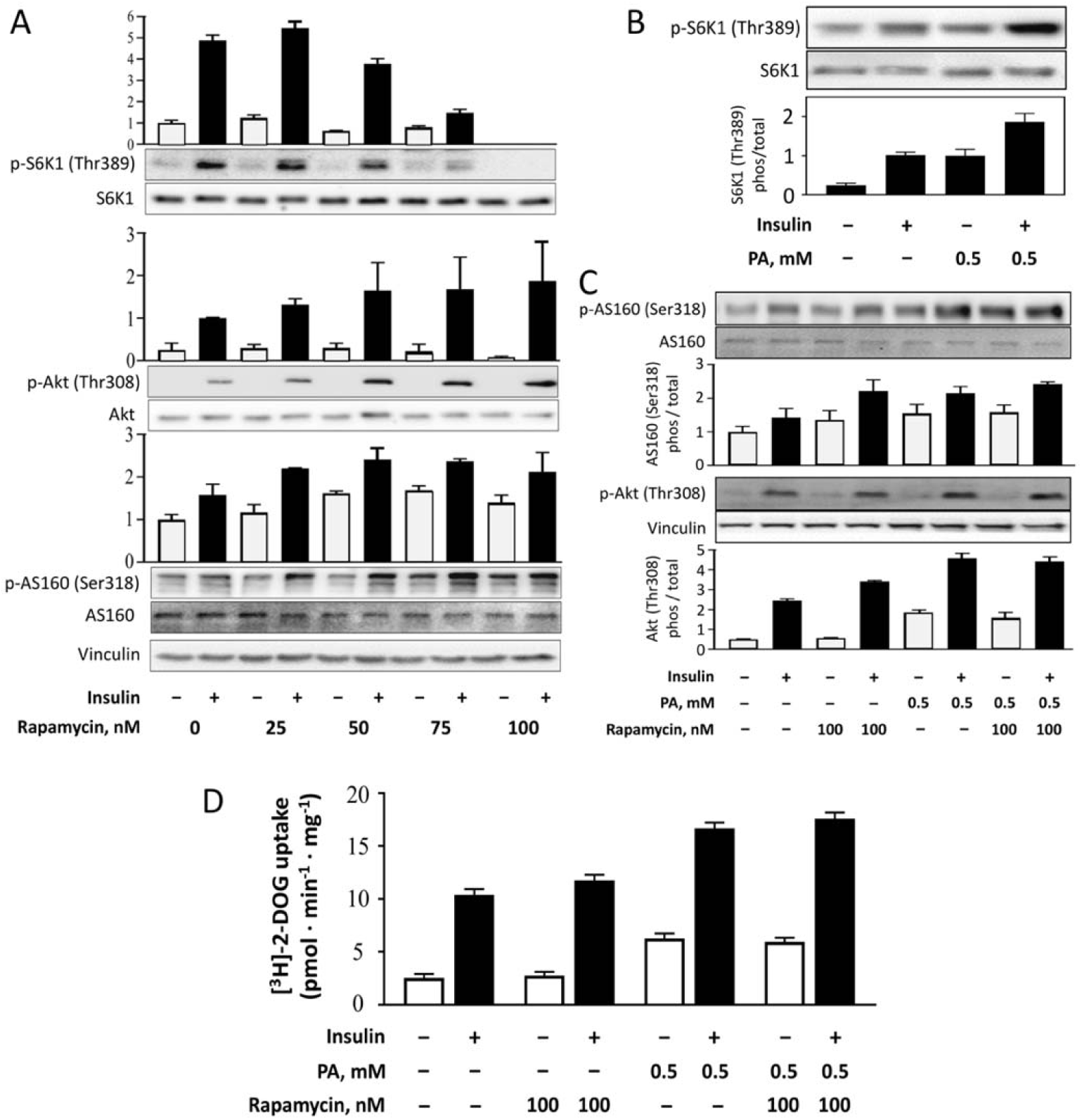
mTORC1-mediated feedback in insulin signaling is not involved in PA-induced increase of basal glucose uptake by 3T3-L1 adipocytes. **(A)** Dose-dependence of rapamycin effects on insulin-induced phosphorylation of S6K1, Akt and AS160 in 3T3-L1 adipocytes assayed immediately after their differentiation in the presence of rosiglitazone. **(B)** PA activates mTORC1 in adipocytes as reported by increased S6K1 phosphorylation in response to cell treatment with 0.5 mM PA for 24 hours. **(C)** PA increases phosphorylation of Akt and AS160 independent of rapamycin treatment of adipocytes. **(D)** PA increases 2-DOG uptake by adipocytes independent of their rapamycin treatment.

To further substantiate the lack of involvement of mTORC1-mediated feedback in PA-driven increase in basal glucose uptake, we performed the inhibitory analysis of [^3^H]-2-DOG uptake by 3T3-L1 adipocytes (Fig. 4D). Similar to the above results (Fig. 3B), 0.5 mM PA increased basal 2-DOG uptake ∼2.5-fold, although reduced its insulin stimulation only ∼2-fold. Notably, these effects of PA were not affected by rapamycin, once again indicating that PA increases basal glucose uptake independent of signaling via mTORC1.

### Excessive palmitate induces insulin resistance in rosiglitazone-differentiated 3T3-L1 adipocytes

Contrary to basal glucose uptake and insulin signaling, which were increased by low PA concentrations (Fig. 4C,D), the insulin-stimulated glucose uptake was reduced by 48-hr exposure to high concentration of PA (1 mM) in adipocytes differentiated in the presence of rosiglitazone (Fig. 3B). To assess the effect of this PA concentration on insulin signaling, we treated adipocytes with 1 mM PA for 24 or 48 hours and probed for insulin signaling. As shown in Fig. 5A, 1 mM PA effectively inhibited insulin signaling at both time intervals. The proximal pathway leading to glucose uptake was inhibited almost completely, whereas phosphorylation of IRS-1 at Tyr612, which is critical to PI3-kinase activation, was not fully inhibited. These results raise the possibility that some other branches of insulin signaling network, such as its mitogenic arm or Cbl/TC10 ubiquitin-ligase pathway (Burchfield et al., 2025) may remain partially active to support some responses of cells to insulin. Notably, AMPK was also responsive to high-PA treatment: while its activity was detectable and little, if any, affected by insulin in the untreated cells, it was fully inhibited by 1-mM PA (Fig. 5A). Finally, high-PA did not affect the cellular content of GLUT1 and GLUT4 (Fig. 5B), ruling out the direct PA effects on glucose transporter expression/degradation.

**Fig. 5.**
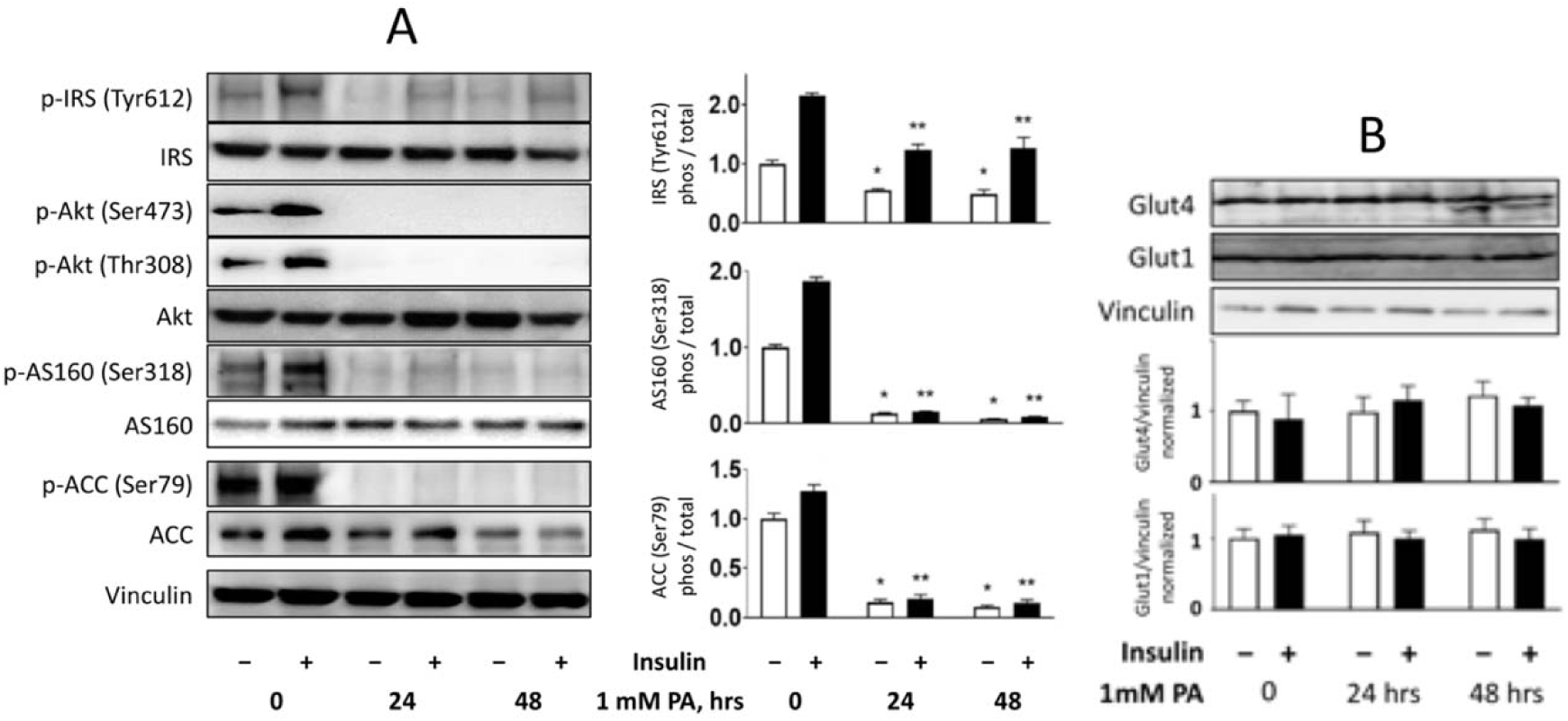
Insulin signaling to glucose uptake is abolished by high PA concentrations in 3T3-L1 adipocytes. **(A)** Representative western blots (*left*) and their statistical quantitation (*right*) of adipocytes treated for 48 hours by 1 mM PA-BSA. **(B)** PA treatment does not affect GLUT1 or GLUT4 content in adipocytes. Data are shown as means ± SD (n=2); *, p <0.05 and **, p <0.01 compared to values in the absence of PA.

Altogether, the clear dissociation between fully inhibited insulin signalling (Fig. 5A) and increased basal glucose uptake (Fig. 3B) suggests that the mechanism underlying the effect of high PA concentration on basal glucose uptake is independent of its effect on insulin signaling along its branch to glucose uptake.

### Increased basal glucose uptake is due to increased GLUT4 exposure at the plasma membrane

To investigate whether the effects of PA on basal glucose uptake are mediated by increased exposure of glucose transporters, we transduced 3T3-L1 preadipocyte with lentiviral construct expressing cMyc-Glut4-nCherry chimera (Fig. 6A). Using mCherry fluorescence as a readout, we sorted out the population of cells with medium level of the chimera expression, propagated the cells and differentiated them into immature adipocytes without rosiglitazone treatment. We additionally cultured resulting adipocytes for a week in order cells to develop larger lipid droplets, but retain insulin stimulation of glucose uptake (cf. Fig. 2D). These adipocyte cultures displayed dense appearance and comparable, though not even, mCherry expression in individual cells (Fig. 6B).

**Fig. 6.**
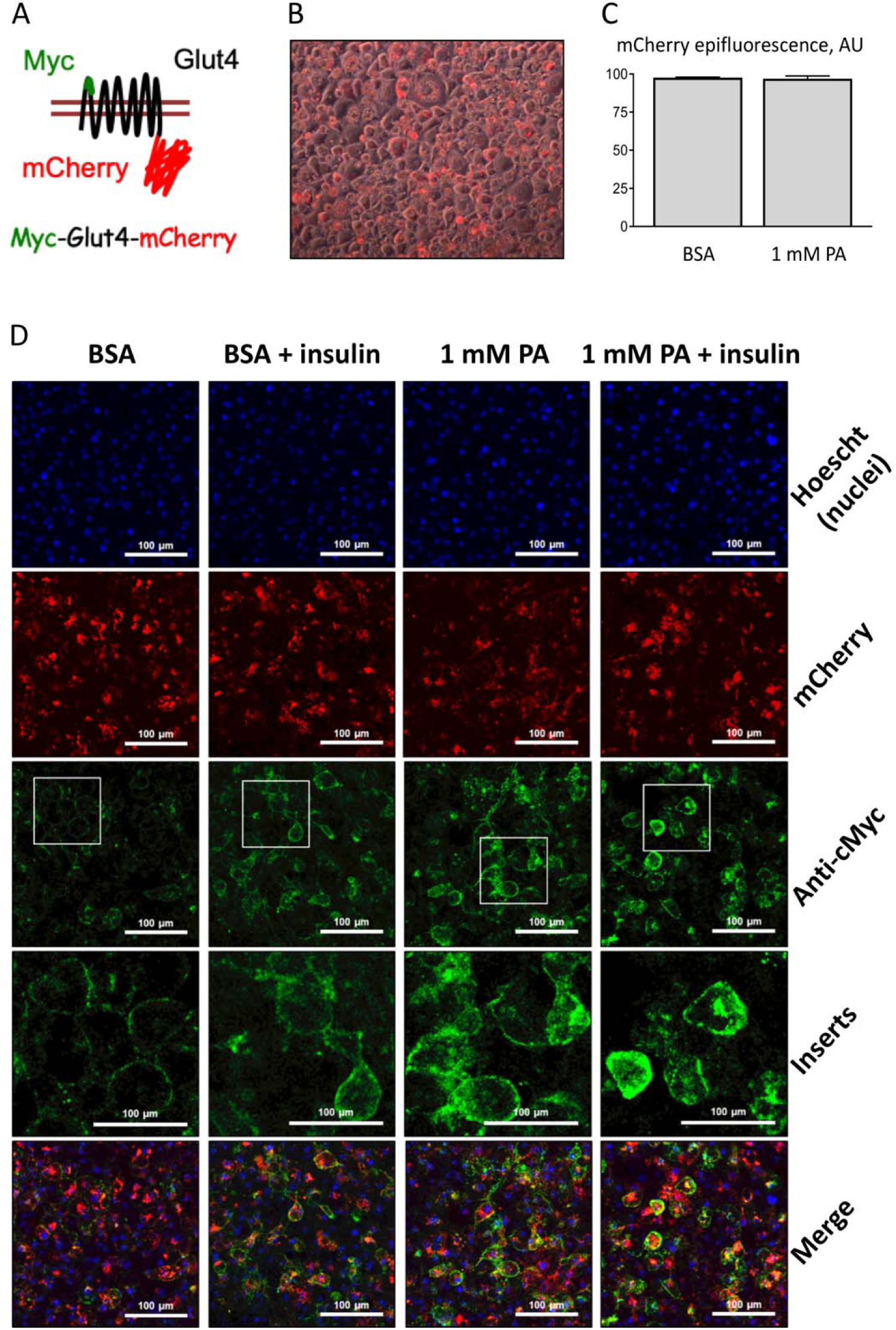
PA increases insulin-independent GLUT4 exposure at the adipocyte plasma membrane. (A) Cartoon illustrating composition of cMyc-Glut4-nCherry chimera. While the C-terminal mCherry tag always resides in the intracellular space and reflects total chimera content, the cMyc tag is located inside the N-terminal extracellular loop of the transporter and becomes accessible to the cognate antibody when the transporter is exposed at the plasma membrane. (B) 3T3-L1 adipocytes that have been differentiated from preadipocytes treated by lentivirus encoding the chimera expressing plasmid and sorted for medium level of mCherry fluorescence. (C) PA treatment for 48 hours of 3T3-L1 adipocytes expressing the cMyc-Glut4-nCherry chimera does not alter their GLUT1 and GLUT4 content. (D) Confocal images of 3T3-L1 adipocytes expressing cMyc-Glut4-nCherry chimera and treated by BSA, or 1 mM PA-BSA for 48 hours, or insulin for 30 min as indicated. For annotation of fluorescence signals refer to the legend on the right.

3T3-L1 adipocytes expressing cMyc-Glut4-nCherry chimera were treated for 48 hours with either BSA alone or 1 mM PA complexed to BSA as above. Using mCherry fluorescence as a readout, we established that PA treatment did not alter the chimera content in these adipocytes (Fig. 6C). The adipocytes were stimulated or not with insulin, fixed and processed for confocal image analysis of Glut4 exposure at the plasma membrane. The analysis was performed in parallel fashion using same settings to ensure that the comparable florescence signals are acquired from all cells. Insulin induced translocation and exposure of the Glut4 chimera at the plasma membrane in the PA-untreated adipocytes, evident by increased green fluorescence of anti-cMyc antibody (Fig. 6D, anti-cMyc). In contrast, PA-treated adipocytes revealed no difference in anti-cMyc fluorescence that was equally increased in unstimulated and insulin-stimulated cells. Notably the anti-cMyc signal appeared mostly as thin rings around the cells, indicative of its plasma membrane location (Fig. 6D, inserts). Imaging the cell nuclei and mCherry fluorescence further indicated that the number of cells and their chimera content were comparable. These results demonstrate that PA-induced basal glucose uptake is mediated by increased Glut4 exposure at the plasma membrane.

## DISCUSSION

In the present study, we investigated the effects of exogenous FFA, primarily PA, on glucose uptake in 3T3-L1 adipocytes, focusing on the role of adipocyte maturation and differentiation conditions. Our findings reveal a complex, context-dependent action of PA that challenges the simplified view of saturated FFA as universal inducers of insulin resistance (Petersen, Shulman, 2018, Samuel et al., 2010), at least for adipocytes with regard to glucose consumption and its intracellular fate.

The key observation is that in commonly used and conventionally differentiated 3T3-L1 adipocytes FFA had no acute effects on lipid accumulation and glucose uptake, however adipocytes increased glucose uptake during long-term maturation in culture (Fig. 1). Notably, the basal glucose uptake increased more than insulin-stimulated, levelling up to insulin-stimulated uptake. This is consistent with observations that adipocytes mostly rely on glucose provision to support their metabolism and increase their fat stores (Krycer et al., 2020, Krycer et al., 2020). In cultured cells this is likely supported for by continuous presence of high glucose in culture medium.

The inherent distinction of fat cell metabolism from that in other cells (Morigny et al., 2021) may underlie fundamental differences in their metabolism extending beyond the common assumption of substrate competition originally described for muscle cells or hepatocytes (Hebbachi, Saggerson, 2012, Randle et al., 1963, McGarry 2002). In addition, as cultured cells are commonly used to model the in vivo behavior, the day-to-day rate of metabolic activity has to be considered as it likely results in fluctuations in nutrient and micronutrient availability, which in turn may result in dynamic changes in cellular metabolism (Tan et al., 2025). This highlights the critical importance of defining the adipocyte model when studying nutrient signaling and metabolism, supporting the notion that availability of essentials engaged by cells from the cell media is an important factor to tune metabolism (Vorotnikov et al., 2022). The overall rate of glucose utilization by adipocyte, which may affect the rate of adipocyte maturation and concurrent metabolic reprogramming, was not directly assessed in our study and remains for the future studies.

We observed that mature adipocytes have altered expression of critical markers indicative of intracellular routing and fatty acid metabolism. Long-term culture in high glucose led to a unilocular, “mature” adipocyte phenotype associated with a dramatic increase in fat content, and a loss of fatty acid oxidation markers, CPT1B and UCP-1 (Fig. 2). This metabolic shift towards lipid-storing phenotype likely represents an adaptive response to high glucose availability. Whereas CPT-1 is not commonly used as a marker of white-to-beige phenotype conversion, UCP-1 is widely considered as a marker of brown and beige phenotype (Rosen, Spiegelman, 2014). Thus, it remains for the future studies whether the adipocyte phenotype conversion may be a consequence of metabolic shift driven by high nutrient availability rather than solely by distinct differentiation programs.

Another key and most striking observation is the dissociation between the effects of PA on basal and insulin-stimulated glucose uptake. In rosiglitazone-differentiated adipocytes, PA dose-dependently increased basal glucose uptake, while at high concentrations it suppressed the insulin-stimulated uptake. This dual effect provides a potential explanation for the long-standing contradictions in the literature regarding FFA action on adipocyte glucose metabolism. The outcome likely depends on the experimental model (differentiation protocol, maturity stage), FFA concentration, and exposure time, which may collectively determine the balance between these opposing pathways.

Rosiglitazone is a PPARγ agonist of thiazolidinedione drugs that upregulate lipid metabolism, including triglyceride storage, fatty acid oxidation, adipose tissue remodeling and adipocyte brightening (Lee et al., 2019). Notably, adipocytes newly differentiated with rosiglitazone, had similar CPT-1 and UCP-1 levels and glucose uptake to adipocytes differentiated in its absence (Fig. 3). However, the rosiglitazone-treated cells acquired sensitivity to PA treatment by selectively increasing basal glucose uptake. Notably, similar effect of PA was observed in human adipocytes differentiated from ADSC in the presence of rosiglitazone. This suggests that the sensitivity to PA may be a feature of beige-like adipocyte state, possibly primed by rosiglitazone activation of PPARγ. This may be a result of metabolic reprograming, such as an increase in FFA storage in triglycerides, which augmented turnover increases demand for glucose influx to generate glyceride glycerol (Bally et al., 1960, Hebbachi, Saggerson, 2012). As mature adipocytes display high basal glucose uptake yet little expression of CPT-1 and UCP-1, their lipid metabolism is likely recast to fat storage rather than oxidation. Whether this metabolic conversion is an adaptive response to escape the increased pressure from external fat availability, and whether the beige-like immature adipocytes are particularly vulnerable to excessive FFA and insulin resistance remains to be studied. The observation that mature adipocytes withstand high FFA load (Fig. 1E), whereas the immature rosiglitazone-treated cells develop insulin-resistant state in response to the same high-dose PA treatment (Fig. 3B) argues for this possibility. Further studies are needed to explore these hypotheses.

Our results further suggest that the mechanism underlying PA-induced basal glucose uptake is fundamentally different from the canonical insulin signaling cascade. First, the increase in basal uptake persisted (Fig. 4D) even when insulin signaling to Akt and AS160 was completely inhibited by high-dose PA treatment in rosiglitazone-treated adipocytes (Fig. 5A). Second, this PA effect was independent of the mTORC1-mediated feedback loop, an established modulator of insulin sensitivity, as rapamycin failed to inhibit it (Fig. 4C). Third, PA treatment led to increased exposure of cMyc-GLUT4-mCherry on the plasma membrane in the absence of insulin stimulation. This insulin-independent translocation of GLUT4 is the likely cellular basis for the enhanced basal glucose uptake. However, it is also possible that the increase in basal glucose uptake induced by PA in rosiglitazone-untreated adipocytes does depend on increased insulin signaling (Fig. 4C). Whether this difference is due to rosiglitazone effects on lipid metabolism and/or adipocyte phenotype is currently unclear and has to be studied further.

These findings suggest that saturated FFA may affect GLUT4 trafficking or other related mechanisms through a novel, insulin-independent pathway in adipocytes. This stands in contrast to its established role in skeletal muscle, where saturated FFA primarily impair insulin-dependent GLUT4 translocation (see references in Introduction). The molecular machinery driving the FFA-induced GLUT4 exposure remains to be fully elucidated. Potential candidates include AMPK, which activation has been linked to increased glucose uptake in some contexts. However, AMPK was inhibited in rosiglitazone-treated adipocytes by 1-mM PA (Fig. 5A), yet these cells retained basal glucose uptake. The function of AMPK in adipocytes is still insufficiently understood (Ceddia 2013, Hebbachi, Saggerson, 2012) and deserves an appraisal.

The induction of insulin resistance by high-dose PA in our model aligns with aspects of the classical lipotoxicity paradigm (Petersen, Shulman, 2018, Samuel et al., 2010). Thus, adipocytes may harbor both the classical pathway for FFA-induced insulin resistance and a parallel, compensatory pathway for insulin-independent glucose uptake stimulation. From a physiological perspective, the latter could be viewed as a salvage mechanism. In states of lipid overload and developing systemic insulin resistance, such a mechanism in adipocytes would help direct glucose towards glycerol-3-phosphate synthesis, facilitating the (re)esterification and safe storage of excess FFAs as triglycerides within the adipose tissue depot. This could initially serve a protective function, preventing ectopic lipid accumulation. Its failure or overwhelming by prolonged, severe lipotoxicity might contribute to adipose tissue dysfunction.

In conclusion, our study demonstrates that palmitate exerts two distinct effects on adipocyte glucose metabolism: (1) an insulin-independent stimulation of basal glucose uptake via increased GLUT4 surface exposure, and (2) at higher doses, an inhibition of insulin-stimulated glucose uptake. The manifestation and balance of these effects depend on differentiation history and metabolic state of adipocytes. These findings underscore that the role of saturated fatty acids in metabolic regulation is cell type- and context-specific. They reveal a previously underappreciated adaptive capacity of adipocytes to couple lipid sensing directly to glucose consumption and metabolism, a pathway that may be relevant to coordination of lipid and glucose metabolism.

## ACKNOWLEDGEMENTS

This work was supported by RSF grant #23-75-00027 (https://rscf.ru/project/23-75-00027/).

